# GBRAP: a tool to retrieve, parse and analyze GenBank files of viral and bacterial species

**DOI:** 10.1101/2021.09.21.461110

**Authors:** Chiara Vischioni, Valerio Giaccone, Paolo Catellani, Leonardo Alberghini, Riccardo Miotti Scapin, Cristian Taccioli

## Abstract

**Summary:** GenBank files contain genomic data of sequenced living organisms. Here, we present GBRAP (GenBank Retrieving, Analyzing and Parsing software), a tool written in Python 3 that can be used to easily download, parse and analyze viral and bacterial GenBank files, even when contain more than one genomic sequence for each species. GBRAP can analyze more files simultaneously through single command-line parameters that give as output a single table showing the genomic characteristics of each organism. It is also able to calculate Shannon, LZSS (Lempel–Ziv–Storer–Szymanski) and topological entropy for both the entire genome and its constitutive elements such as genes, rRNAs, tRNAs, tmRNAs and ncRNAs together with Chargaff’s second parity rule scores obtained using different mathematical methods. Moreover, GBRAP can calculate, the number, the length and the nucleotides abundance of genomic components for each DNA strand and for the overlapping regions among the two complementary helixes. To our knowledge, this is the only software capable of providing this type of genomic analyses all together in a single tool, that, therefore can be used by the scientists interested in both genomics and evolutionary research.

**Availability and implementation:** The data underlying this article are available from the corresponding author on reasonable request.

## Introduction

Next Generation Sequencing (NGS) has made possible the sequencing of thousands of genomes in short time, allowing the researchers to download and analyze their genomic information both in the fields of biology and medicine. Nowadays, GenBank files (extension: GBFF) can be download from the National Center for Biotechnology Information (NCBI) web site (https://www.ncbi.nlm.nih.gov/genome) or through the RefSeq and GenBank FTP servers (ftp://ftp.ncbi.nlm.nih.gov/genomes/genbank/ or ftp://ftp.ncbi.nlm.nih.gov/genomes/refseq/). GenBank and RefSeq databases are extensive publicly collections of available nucleotides of more than 400,000 formally described species, ranging from the smallest viruses and plasmids, up to the biggest bacteria. With the aim of simplifying the procedure of downloading one or multiple GBFFs and making possible the contemporary analysis of the information they contain, we developed GBRAP. Our tool is written in Python 3 and is usable as command line through two scripts named gbr.py and gbap.py. The former allows the users to download specific microorganism species, whereas the latter provides a wide range of genomic statistics analyses such as the count, the frequencies and the length of coding regions (CDS), ribosomal RNAs (rRNAs), transfer RNAs (tRNAs), transfer-messenger RNAs (tmRNA) and non-coding RNAs (ncRNAs). Moreover, it counts the nucleotides of regions that overlaps one with another both on the same strand or within them. It also performs, Shannon (Shannon, 1948), LZSS (Storer, 1982) and topological entropy (Koslicki, 2011) and two different genomic Chargaff’s second parity rule (Fariselli, 2020) scores (see Supplementary information for more details about entropy and Chargaff’s rules within genomic sequences). GBRAP is very easy to use and does not require powerful computers, allowing the users to analyze viral and prokaryotic genomes using normal machines. To our knowledge, this is the only tool available (see Supplementary information for software comparison) that offers the features described above, all included in a single software free of charge, publicly available, and easy to use.

## Material and Methods

GBRAP is a unique tool able to download and import GenBank files containing one or more sequences and outputs genomics statistics analyses. It is a command line and user-friendly software and can be installed in all the Operative Systems, without any other package needed, with the exception of Biopython module (see Supplementary information for information about installation). GBRAP is a package containing two Python scripts: the first can be used to download GenBank files (gbr.py), whereas the other one (gbap.py) outputs the analyses in a tab delimited text file that can be easily opened using Excel or any another spreadsheet software. In particular, the RETRIEVE class contained in gbr.py allows the user to download GenBank files from GenBank or RefSeq repositories within the FTP (Files Transfer Protocol) NCBI database. Setting the different parameters, it is possible to choose the species of interest (`-s`), the preferred database (`-d`), the selected taxon (`-t`), the assembly type (`-a`), and the output directory (`-o`). Multiple species can be downloaded at once using regular expression wildcards or retrieving the entire NCBI genome directories such as bacteria, archaea or viral genomes (see Supplementary information). As a result, a single or multiple GBFF files can be downloaded in the chosen directory and used for the subsequent analyses. Once the user has downloaded GBFF files using gbr.py and decompressed them, gbap.py can be used to generate a tab delimited file that includes the genomic features (n=132) processed and analyzed through the ANPA class (ANalysis and Parsing tool). Single or multiple analyses can be chosen using `-i` and `-d` parameters respectively, whereas `-f` can be invoked in order to select the genomic sequences of interest marked by a specific NCBI ID that in GBFF files are named as “LOCUS” (see Supplementary information for details).

## Results and Discussion

In order to train GBRAP, we downloaded the GenBank files of some of the most studied bacterial species in the field of human health and food science: *Clostridium botulinum, Listeria monocytogenes, Escherichia coli, Salmonella enterica, and Yersinia pestis*. According to the analysis output, the genome size of the Firmicutes species analyzed (*Clostridium botulinum and Listeria monocytogenes*) is significantly lower (32%) compared to the one of the examined Proteobacteria (*Escherichia coli, Salmonella enterica, and Yersinia pestis*). Furthermore, our results also showed that the number of genes in *C. Botulinum* and *L. monocytogenes* is lower than that of *E. coli*, *S. enterica* and *Y. pestis*, which is a logical consequence of the fact that bacteria genomes, in general, contain almost exclusively protein-coding sequences (see Supplementary Information). Thanks to the gbap.py script and the ANPA class, we also investigated the degree of genomic information and structural genome stability of the analyzed species. To this aim, we found that *C. Botulinum* and *L. monocytogenes* have a lower amount of genomic information within their genome (Shannon entropy score = 1.90) compared to *E. coli, S. enterica, and Y. pestis* (Shannon entropy score =1.99) and a lower level of genomic stability (0.9 vs 0.99) when calculation Chargaff’s second parity rule scores (see Supplementary information and Figure 1S for details). The same results were obtained by calculating the topological entropy and the LZSS compression score using the aforementioned genomes (see Supplementary information). In this context, looking exclusively to our results, we can state that *C. Botulinum* and *L. monocytogenes* have a lower degree of information abundance and genomic stability compared to the Proteobacteria species examined, underling a lower genetic complexity in the latter class.

## Conclusion

DNA base composition regularities are highly desirable for evolutionary and genomics research efforts. Our tool, in addition to an easy way of downloading GenBank files from NCBI, integrates in a single analysis many features in term of calculation of mathematical DNA and RNA proprieties such as the genome size, the frequencies of base pairs or the entropy energy levels, all included, for the first time in a unique software. In a structural and evolutionary point of view, this integration could be very useful in order to link the sequence regularities with the dynamism of the DNA double helix. Finally, GBRAP can process the genome of any old and newly sequenced organism, giving the researchers the opportunity to analyze microorganism genomes (viruses, bacteria, and archaea species) in a reasonable time and using normal computers.

## Funds

This study has been funded by MAPS Department funds BIRD2021 - prot. BIRD213010, University of Padova.

## Acknowledgements

We would like to thank Professor Alessio Boattini and Doctor Tania Bobbo for the precious advices regarding the Python code used for the ANPA class.

