## Supplementary information for "GBRAP: a tool to retrieve, parse and analyze GenBank files of viral and bacterial species"

[*]To whom the correspondence should be addressed

#### GBRAP Overview

GBRAP (GenBank tool for Retrieving, Analyzing and Parsing software), is a tool written in Python 3 that the researchers can use to download microorganism GenBank files (GBFF) from the FTP servers of the NCBI database. Using GBRAP is also possible to calculate Chargaff's scores, Shannon, LZSS (Lempel–Ziv–Storer–Szymanski), and topological entropies for both the entire genome and its constitutive elements such as genes, rRNAs, tRNAs, tmRNAs and ncRNAs. Moreover, GBRAP is able to calculate the number, the length and the nucleotides abundance of these genomic components for each strand and for the overlapping regions among the two DNA helics.

#### Installation

GBRAP is an open-source tool. Once is downloaded it can be used for downloading GBFF files through the command `python3 ./gbr.py [parameters]` or by typing `python3 ./gbap.py [parameters] > output.txt` when the researchers need to analyze GBFF files. GBRAP includes all the needed files for the execution of the complete analysis, with the exception of the Biopython module. For Ubuntu, Debian, Centos, and Fedora users who do not have "rsync" already present in their system, we suggest to install it. For Windows 10 users, we suggest to install the Ubuntu terminal directly on their system using this Microsoft guide (https://docs.microsoft.com/en-us/windows/wsl/install-win10) and then install Biopython module and rsync.

#### Usage

GBRAP is made of two scripts. Gbr.py is aimed to download GenBank files from RefSeq and GenBank repositories, whereas gbap.py is aimed to analyze the GBFF files retrieved from RefSeq and GenBank databases. In particular, gbr.py requires 5 mandatory fixed parameters (`-s`, `-d`, `-t`, `-a`, and `-o`), which can be modified accordingly to the user specific needs (see more information below). If one of them is not specified, the connection with the NCBI server will be blocked, otherwise the download will start immediately. It is also possible to download the entire contents of a specific directory within the database. Gbap.py instead requires only two parameters `-i` (input file) and `-o` (name of the directory where the input file is located). It is important to remember that GBFF files should be decompressed before being analyzed.

### Downloading GenBank files script

Gbr.py contains a class (RETRIEVE) that allows the user to download GenBank files (*.gbff) from GenBank or RefSeq directories within NCBI database. The output files will be downloaded in the directory chosen using the `-o` parameter (see below for details). This is a brief description of the commands:

- [-s], [--species]: e.g. -s Yersinia_pestis or e.g. -s Yersinia*

This parameter is used to select the name of the organism to be downloaded. In order to avoid errors, the white space between genus and species has to be replaced by ‘’_’’. If the underscore is missing, the program will give an “unrecognize argument error”, because the attribute will be seen as a command. The user can also obtain all the genomes included in the same genus, replacing the name of the species with a “*” (note that the double quotes are mandatory in some operative system such as Mac OS).

- [-d], [--database]: e.g. -d genbank

This parameter opts for GenBank database or RefSeq one.

- [-t], [--taxon]: e.g. -t bacteria

This parameter discriminates between classes or reigns of organisms in which the genome call will be performed. The user should write one among 3 different choices: archaea, bacteria, and viral.

- [-a], [--assembly]: e.g. -a reference

This parameter allows the user to choose within different types of assembly versions (latest_assembly_versions, representative, reference, and all_assembly_versions). It is also possible to download both the reference and the representative using the command re*.

- [-o], [--outputpath]: e.g. -o /home/user/Desktop/

The user has to set up this parameter to determine the path of the downloading. It is also important to check if the selected directory exists or not.

To summarize, the RETRIEVE class included within the gbr.py script, allows the users to download gbff files from GenBank and RefSeq repositories.

#### Usage example

If a user wants to download the last assembly version of the genome of *Yersinia pestis* from the FTP RefSeq repository in the directory called “my_directory”, the command to run is:

```
python3 ./gbr.py -s Yersinia_pestis -d refseq -t bacteria -a latest_assembly_versions -o /my_directory
```

As result the user will obtain in the selected directory a compressed file (.gbff.gz) containing the genomic sequence of interest.

### Analysis of GenBank files tool

The gbap.py script works through the class ANPA and allows the users to parse and perform genomic analyses using all the features contained in GenBank files (.gbff). The files need to be uncompressed before being used. This is a brief description of the commands:

- [-i], [--input]: e.g. -i xxx_genomic.gbff

This parameter is used to select a specific gbff file.

- [-d], [--directory]: e.g. -d home/user/Desktop/downloaded_genomes/

This parameter allows the analysis of all the genomic gbff files within a specific directory.

- [-f], [--filter]: e.g. -f NC_00

The user can set this parameter in order to filter out the LOCUS not needed for a specific analysis. For example, the user might remove of the LOCUS starting with NC_00.

#### Usage example

To summarize the ANPA class employment, here we report an example of its usage. The aim of the test is to remove all the genomes starting the string NC_XXX:

./gbap.py -i xxx_genomic.gbff -f NC_XXX > output.txt

or, if there are multiple gbff files in the folder the script should be like:

./gbap.py -d home/user/Desktop/downloaded_genomes/ -f NC_XXX > output.txt

ANPA returns in output a tab delimited text file that could be imported in excel. Each file contains:

- ID: GenBank identification code;
- name: organism's name;
- taxon: organism taxon belonging;
- bp_genome: total number of bases in the genomic sequence;
- bp_genA: total number of Adenines in the genomic sequence;
- bp_genT: total number of Thymines in the genomic sequence;
- bp_genC: total number of Cytosines in the genomic sequence;
- bp_genG: total number of Guanines in the genomic sequence;
- fr_genA: frequency of Adenine nucleotides in the genomic sequence;
- fr_genT: frequency of Thymine nucleotides in the genomic sequence;
- fr_genC: frequency of Cytosine nucleotides in the genomic sequence;
- fr_genG: frequency of Guanine nucleotides in the genomic sequence;

- genomic_topological_entropy_score: Topological score calculated on genomic sequences using Koslicki method (Koslicki et al. 2011);
- genomic_chargaff_score_pf: Chargaff's second parity rule score calculated on genomic sequences as shown below*;
- genomic_chargaff_score_ct: Chargaff score calculated on genomic sequences using CT method as shown below**;
- genomic_shannon_score: Shannon*** score calculated on genomic sequences using Shannon method (Shannon, 1948);
- genomic_vLZSS_score: LZSS score calculated on genomic sequences as shown below****;
- bp_cds_total: total number of bases in the coding sequences;
- bp_cdsA: total number of Adenines in the coding sequences;
- bp_cdsT: total number of Thymines in the coding sequences;
- bp_cdsC: total number of Cytosines in the coding sequences;
- bp_cdsG: total number of Guanines in the coding sequences;
- fr_cdsA: frequency of Adenine nucleobase in the coding sequences;
- fr_cdsT: frequency of Thymine nucleobase in the coding sequences;
- fr_cdsC: frequency of Cytosine nucleobase in the coding sequences;
- fr_cdsG: frequency of Guanine nucleobase in the coding sequences;
- bp_cds_overlap_plus: number of bases overlapping the + strand of the coding sequences;
- bp_cds_overlap_minus: number of bases overlapping the - strand of the coding sequences;
- bp_cds_overlap_plus_vs_minus: number of bases overlapping + and – strands of the coding sequences;
- n_cds_overlaps_plus: number of CDSs overlapping the + strand of the genomic sequence;
- n_cds_overlaps_minus: number of CDSs overlapping the - strand of the genomic sequence;
- n_cds_overlaps_total: number of CDSs overlapping the two strands of the genomic sequence;
- n_cds_plus: number of coding sequences on the + strand of the genomic sequence;
- n_cds_minus: number of coding sequences on the - strand of the genomic sequence;
- n_cds_total: total number of coding sequences in the genomic sequence;
- cds_topological_entropy_score: Topological score† calculated on cds sequences using Koslicki method (Koslicki, 2011);
- cds_chargaff_score_pf: Chargaff's second parity rule score calculated on cds sequences as shown below*;
- cds_chargaff_score_ct: Chargaff score calculated on cds sequences using CT method as shown below**;
- cds_shannon_score: Shannon*** score calculated on cds sequences using Shannon method (Shannon, 1948);
- cds_vLZSS_score: LZSS score (Storer et al., 1982) calculated on cds sequences as shown below****;
- bp_rRNA_total: total number of bases in the rRNA sequences;
- bp_rRNA_A: total number of Adenines in the rRNA sequences;
- bp_rRNA_T: total number of Thymines in the rRNA sequences;
- bp_rRNA_C: total number of Cytosines in the rRNA sequences;
- bp_rRNA_G: total number of Guanines in the rRNA sequences;
- fr_rRNA_A: frequency of Adenine nucleobase in the rRNA sequences;

- fr_rRNA_T: frequency of Thymine nucleobase in the rRNA sequences;
- fr_rRNA_C: frequency of Cytosine nucleobase in the rRNA sequences;
- fr_rRNA_G: frequency of Guanine nucleobase in the rRNA sequences;
- bp_rRNA_overlap_plus: number of bases overlapping the + strand of the rRNA sequences;
- bp_rRNA_overlap_minus: number of bases overlapping the - strand of the rRNA sequences;
- bp_rRNA_overlap_plus_vs_minus: number of bases overlapping + and – strands of the rRNA sequences;
- n_rRNA_overlaps_plus: number of rRNAs overlapping the + strand of the genomic sequence;
- n_rRNA_overlaps_minus: number of rRNAs overlapping the - strand of the genomic sequence;
- n_rRNA_overlaps_total: number of rRNAs overlapping the two strands of the genomic sequence;
- n_rRNA_plus: number of rRNAs on the + strand of the genomic sequence;
- n_rRNA_minus: number of rRNAs on the - strand of the genomic sequence;
- n_rRNA_total: total number of rRNAs in the genomic sequence;
- rRNA_topological_entropy_score: Topological score† calculated on rRNA sequences using Koslicki method (Koslicki, 2011);
- rRNA_chargaff_score_pf: Chargaff's second parity rule score calculated on rRNA sequences as shown below*;
- rRNA_chargaff_score_ct: Chargaff score calculated on rRNA sequences using CT method as shown below**;
- rRNA_shannon_score: Shannon*** score calculated on rRNA sequences using Shannon method (Shannon, 1948);
- rRNA_vLZSS_score: LZSS score (Storer et al., 1982) score calculated on rRNA sequences as shown below****;
- bp_tRNA_total: total number of bases in the tRNA sequences;
- bp_tRNA_A: total number of Adenines in the tRNA sequences;
- bp_tRNA_T: total number of Thymines in the tRNA sequences;
- bp_tRNA_C: total number of Cytosines in the tRNA sequences;
- bp_tRNA_G: total number of Guanines in the tRNA sequences;
- fr_tRNA_A: frequency of Adenine nucleobase in the tRNA sequences;
- fr_tRNA_T: frequency of Thymine nucleobase in the tRNA sequences;
- fr_tRNA_C: frequency of Cytosine nucleobase in the tRNA sequences;
- fr_tRNA_G: frequency of Guanine nucleobase in the tRNA sequences;
- bp_tRNA_overlap_plus: number of bases overlapping the + strand of the tRNA sequences;
- bp_tRNA_overlap_minus: number of bases overlapping the - strand of the tRNA sequences;
- bp_tRNA_overlap_plus_vs_minus: number of bases overlapping + and – strands of the tRNA sequences;
- n_tRNA_overlaps_plus: number of tRNAs overlapping the + strand of the genomic sequence;
- n_tRNA_overlaps_minus: number of tRNAs overlapping the - strand of the genomic sequence;
- n_tRNA_overlaps_total: number of tRNAs overlapping the two strands of the genomic sequence;
- n_tRNA_plus: number of tRNAs on the + strand of the genomic sequence;

- n_tRNA_minus: number of tRNAs on the - strand of the genomic sequence;
- n_tRNA_total: total number of tRNAs in the genomic sequence;
- tRNA_topological_entropy_score: Topological score† calculated on tRNA sequences using Koslicki method (Koslicki et al., 2011);
- tRNA_chargaff_score_pf: Chargaff's second parity rule score calculated on tRNA sequences as shown below*;
- tRNA_chargaff_score_ct: Chargaff score calculated on tRNA sequences using CT method as shown below**;
- tRNA_shannon_score: Shannon*** score calculated on tRNA sequences using Shannon method (Shannon, 1948);
- tRNA_vLZSS_score: LZSS score (Storer et al., 1982) calculated on tRNA sequences as shown below****;
- bp_ncRNA_total: total number of bases in the ncRNA sequences;
- bp_ncRNA_A: total number of Adenines in the ncRNA sequences;
- bp_ncRNA_T: total number of Thymines in the ncRNA sequences;
- bp_ncRNA_C: total number of Cytosines in the ncRNA sequences;
- bp_ncRNA_G: total number of Guanines in the ncRNA sequences;
- fr_ncRNA_A: frequency of Adenine nucleobase in the ncRNA sequences;
- fr_ncRNA_T: frequency of Thymine nucleobase in the ncRNA sequences;
- fr_ncRNA_C: frequency of Cytosine nucleobase in the ncRNA sequences;
- fr_ncRNA_G: frequency of Guanine nucleobase in the ncRNA sequences;
- bp_ncRNA_overlap_plus: number of bases overlapping the + strand of the ncRNA sequences;
- bp_ncRNA_overlap_minus: number of bases overlapping the - strand of the ncRNA sequences;
- bp_ncRNA_overlap_plus_vs_minus: number of bases overlapping + and – strands of the ncRNA sequences;
- n_ncRNA_overlaps_plus: number of ncRNAs overlapping the + strand of the genomic sequence;
- n_ncRNA_overlaps_minus: number of ncRNAs overlapping the - strand of the genomic sequence;
- n_ncRNA_overlaps_total: number of ncRNAs overlapping the two strands of the genomic sequence;
- n_ncRNA_plus: number of ncRNAs on the + strand of the genomic sequence;
- n_ncRNA_minus: number of ncRNAs on the - strand of the genomic sequence;
- n_ncRNA_total: total number of ncRNAs in the genomic sequence;
- ncRNA_topological_entropy_score: Topological score† calculated on ncRNA sequences using Koslicki method (Koslicki et al., 2011);
- ncRNA_chargaff_score_pf: Chargaff's second parity rule score calculated on ncRNA sequences as shown below*;
- ncRNA_chargaff_score_ct: Chargaff score calculated on ncRNA sequences using CT method as shown below**;
- ncRNA_shannon_score: Shannon*** score calculated on ncRNA sequences using Shannon method (Shannon, 1948);
- tmRNA_vLZSS_score: LZSS score (Storer et al., 1982) calculated on tmRNA sequences as shown below****;

- bp_tmRNA_total: total number of bases in the tmRNA sequences;
- bp_tmRNA_A: total number of Adenines in the tmRNA sequences;
- bp_tmRNA_T: total number of Thymines in the tmRNA sequences;
- bp_tmRNA_C: total number of Cytosines in the tmRNA sequences;
- bp_tmRNA_G: total number of Guanines in the tmRNA sequences;
- fr_tmRNA_A: frequency of Adenine nucleobase in the tmRNA sequences;
- fr_tmRNA_T: frequency of Thymine nucleobase in the tmRNA sequences;
- fr_tmRNA_C: frequency of Cytosine nucleobase in the tmRNA sequences;
- fr_tmRNA_G: frequency of Guanine nucleobase in the tmRNA sequences;
- bp_tmRNA_overlap_plus: number of bases overlapping the + strand of the tmRNA sequences;
- bp_tmRNA_overlap_minus: number of bases overlapping the - strand of the tmRNA sequences;
- bp_tmRNA_overlap_plus_vs_minus: number of bases overlapping + and – strands of the tmRNA sequences;
- n_tmRNA_overlaps_plus: number of tmRNAs overlapping the + strand of the genomic sequence;
- n_tmRNA_overlaps_minus: number of tmRNAs overlapping the - strand of the genomic sequence;
- n_tmRNA_overlaps_total: number of tmRNAs overlapping the two strands of the genomic sequence;
- n_tmRNA_plus: number of tmRNAs on the + strand of the genomic sequence;
- n_tmRNA_minus: number of tmRNAs on the - strand of the genomic sequence;
- n_tmRNA_total: total number of tmRNAs in the genomic sequence;
- tmRNA_topological_entropy_score: Topological score† calculated on tmRNA sequences using Koslicki method (Koslicki et al., 2011);
- tmRNA_chargaff_score_pf: Chargaff's second parity rule score calculated on tmRNA sequences as shown below*;
- tmRNA_chargaff_score_ct: Chargaff score calculated on tmRNA sequences using CT method as shown below**;
- tmRNA_shannon_score: Shannon*** score calculated on tmRNA sequences using Shannon method (Shannon, 1948);
- tmRNA_vLZSS_score: LZSS score (Storer et al., 1982) calculated on tmRNA sequences as shown below****.

### Chargaff’s second parity rule meaning

In 1950, Erwin Chargaff discovered that the four nucleotides contained in a DNA double helix (A=Adenine, T=Thymine, C=Cytosine and G=Guanine) are symmetrically abundant in both strands of DNA. This symmetry was called Chargaff's first parity rule. In 1968, Chargaff also discovered that also on each DNA strand, the number of Adenines is almost equal to that of Thymines and the number of Cytosines is almost equal to that of Guanines. The first rule was easily explained by the fact that within DNA strands A matches with T, whereas C matches with G. On the single strand, however, this symmetry (Chargaff's second parity rule) is not easily explained. In 2020 four Italian researchers (Fariselli et al., 2020) discovered that this symmetry is linked to the energy of the DNA molecule which has greater stability when it has a Chargaff's second parity rule score close to one.

##### * Chargaff's second parity rule score calculated using an easy method (PF)

An easy way to calculate Chargaff’s second parity rule is this:

$$\mathrm{ABS}\left(\frac{(\#\mathrm{A}-\#\mathrm{T})}{(\#\mathrm{A}+\#\mathrm{T})} + \frac{(\#\mathrm{C}-\#\mathrm{G})}{(\#\mathrm{C}+\#\mathrm{G})}\right)$$

where "#" means "number of" and A = Adenines, T = Thymines, C = Cytosines and G = Guanines.

In this way the perfect Chargaff’s second parity rule score is zero such as in case of “ATGC”. The minimum value depends on the length of the sequence. Python 3 code is as follows:

```
def Chargaff_PF(self,sequence):
    ''' It counts the number of bases in a sequence'''
    A=sequence.count("A")
    T=sequence.count("T")
    C=sequence.count("C")
    G=sequence.count("G")
    nA=a+A
    nT=t+T
    nC=c+C
    nG=g+G
    return abs((nA-nT)/(nA+nT)) + abs((nC-nG)/(nC+nG))
```

##### ** Chargaff's second parity rule score calculated using Cristian Taccioli (CT) method

Chargaff’s second parity rule score calculated using Cristian Taccioli’s method is:

$$\frac{\left(\frac{\#A}{\#T}+\frac{\#C}{\#G}\right)}{2}, where\ \#T\ and\ \#G\ \neq 0$$

*The bases with the highest value must be placed in the denominator.*
*In this example A and C have a lower value than T and G respectively.*

"#" means "number of" and A = Adenines, T = Thymines, C = Cytosines and G = Guanines.

Using this equation Chargaff’s second parity rule score is always between zero and one, where one is the maximum value. For example, the sequence “ATGC” has a perfect score which is one because the number of A is equal to the number of T, whereas the number of C is equal to the number of G.

This score does not depend on sequence length. Chargaff's second parity rule calculated using C.T method can be calculated in Python 3 as follow:

```
 def count_bases(self,sequence):
    ''' It counts the number of bases in a sequence'''
    A=sequence.count("A")
    T=sequence.count("T")
    C=sequence.count("C")
    G=sequence.count("G")
    a=sequence.count("a")
    t=sequence.count("t")
    c=sequence.count("c")
    g=sequence.count("g")
    nA=a+A
    nT=t+T
    nC=c+C
    nG=g+G
    tot=nA+nT+nC+nG
    return [tot,nA,nT,nC,nG]

def chargaff_CT(s):
    ''' Method for Shannon entropy and Chargaff score by Cristian Taccioli '''
    s = str(s).replace("N","")
    nt = count_bases(s)

    if nt[0] !=0 and nt[1] !=0 and nt[2] !=0 and nt[3] !=0 and nt[4] !=0:
        pA = float(nt[1]/nt[0])
        pT = float(nt[2]/nt[0])
        pC = float(nt[3]/nt[0])
        pG = float(nt[4]/nt[0])

        if nt[1] <= nt[2]:
            rAT = nt[1]/nt[2]
        else:
            rAT = nt[2]/nt[1]
        if nt[3] <= nt[4]:
            rCG = nt[3]/nt[4]
        else:
            rCG = nt[4]/nt[3]
        result_ct = (rAT+rCG)/2
    else:
        try:
            pA = float(nt[1]/nt[0])
        except:
            pA = 'NA'

        try:
            pT = float(nt[2]/nt[0])
        except:
            pT = 'NA'

        try:
            pC = float(nt[3]/nt[0])
        except:
            pC = 'NA'

        try:
            pG = float(nt[4]/nt[0])
        except:
            pG = 'NA'
```

```
        result_ct = 'NA'

chargaff_CT('ATGC')
```

#### Entropy scores calculation

The concept of entropy was introduced in the early 19th century by Rudolf Julius Emanuel Clausius. It represents a characteristic quantity of the state of a physical system capable of expressing the ability of the system itself to be able to proceed to spontaneous transformations and, consequently, the loss of ability to do work when such transformations occur. In simplified terms, the value of entropy increases when the system undergoes spontaneous variations and therefore loses part of its ability to undergo such variations and perform work. In 1872 Ludwig Boltzmann generalized this concept through the study of statistical mechanics by defining entropy as the degree of disorder of a system. In 1948 Claude Elwood Shannon equated the degree of inaccuracy of a message with disorder. For Shannon, in fact, the entropy of information was the degree of complexity of a message that represents the minimum average number of symbols necessary for the encoding of the message itself.

##### *** Shannon score

The term entropy in information sciences was introduced by Shannon in the paper "A Mathematical Theory of Communication" (Shannon, 1948). Shannon entropy is calculated as follows:

$$\boldsymbol{H}(\boldsymbol{X}) = -\sum_{i=1}^{n} \boldsymbol{P}(\boldsymbol{x_i})\boldsymbol{logP}\,(\boldsymbol{x_i}),\ \textit{where P is the frequency of nucleotides}$$

Python 3 code for Shannon entropy is:

```
import math
from collections import Counter

def entropy(s):
''' from rosettacode.org '''
      p, lns = Counter(s), float(len(s))
      return -sum(count/lns * math.log(count/lns, 2) for count in p.values())
```

##### **** Lempel–Ziv–Storer–Szymanski score

LZSS was calculated using the methods published by Storer et al. in 1982 (Storer et al. 1982). LZSS measures the compression of a message and virtually returns its complexity. The more a code is difficult to compress, the more it contains information. In this sense, the higher the LZSS value, the more it is incompressible and therefore it has more information. It can be calculated in Python 3 in this way:

```
def kolmogorov(self,s):
    ''' Calculate DEFLATE compression, that is a variation of LZSS compression
algorithm (Lempel-Ziv-Storer-Szymanski) '''
    l = float(len(s))
    if (l > 0) or (l != None):
        compr = zlib.compress(s)
        c = float(len(compr))

        try:
            result = c/l
        except:
```

```
            result = 'NA'
    else:
        result = 'NA'
        return result
```

† Topological entropy score

Topological entropy is a non-negative real number that is capable of measuring the complexity of a message. Topological entropy was first introduced in 1965 by Adler, Konheim and McAndrew (Adler et al. 1965). In 2011 Koslicki has defined a new approximation to topological entropy free from the finite sample effects and high-dimensionality problems. The formula and code can be retrieved from Koslicki, 2011 (Koslicki et al. 2011).

### Performance

We tested GBRAP on three computers:

1) An Intel Xeon W-2195 2.30 GHz desktop computer;
2) An Intel Core i7 CPU 8th Gen 2.8 GHz laptop;
3) An Intel Centrino processor 1.6 GHz laptop.

These machines have 126, 16 and 1 GB of RAM respectively. The first computer performed the downloading and analysis of *Escherichia coli* K-12 substr. MG1655, (GCA_000005845.2_ASM584v2_genomic.gbff) in 1 minutes and 52 seconds using 0.1% of RAM (maximum value), the second machine performed in 5 minutes and 31 seconds using 2.6% of RAM in average (maximum value), while the last one performed the analysis in 12 minutes and 42 seconds using 11% (maximum value) of RAM. All these times were calculated performing the tests a hundred times and the values presented here are the median of all the test values.

### Comparisons with other software

To our knowledge, three different tools are available to parse GenBank files. However, none of them is able to analyze GenBank files features in the way GBRAP does. In particular GBRAP is the only tool able to both download and give in output more than 130 genomic variables calculated as Chargaff's second parity rule scores, different types of entropy values and a specific set of calculations for each genomic element such as genes, rRNAs, tRNAs, tmRNAs, and ncRNAs. This is a list of software similar to GBRAP:

- **Genbankr** (Becker et al., 2019) is an R package, whose basic options consisting in functions to parse GenBank files and displaying annotations of the genomic elements within a species.
- **GBParsy/GBParsyPy** (Lee et al., 2008) is a C-language and Python based library-catalogue that includes specific functions built with the aim of parsing GenBank files, giving back as output data, reference, feature, location and qualifier information. Furthermore, GBParsyPy is a specific Python version of the same tool, that gets all the features from the previous GBParsy.
- **GeneRecords** (D'Addabbo et al., 2004) is a FileMaker pro engine-based tool, built as storage and retrieval system of genetic data, useful to parse and manipulate GenBank flat files and extract desired data from a database entry.

### Analysis examples

GenBank files hold more information than FASTA format files, and for this reason are much more difficult to parse when trying to retrieve useful information. Table1S shows just 8 of the 132 features that can be retrieved and further investigated using gbap.py.

| NCBI Id | Species | Class | Size | fr_genA | fr_genT | fr_genC | fr_genG | n_cds |
|---|---|---|---|---|---|---|---|---|
| **CP000727.1** | *Clostridium botulinum* | Firmicutes | 3,760,560 | 0.35 | 0.36 | 0.14 | 0.14 | 3403 |
| **AL591824.1** | *Listeria monocytogenes* | Firmicutes | 2,944,528 | 0.31 | 0.31 | 0.19 | 0.19 | 2855 |
| **CP003289.1** | *Escherichia coli* | Proteobacteria | 5,273,097 | 0.25 | 0.25 | 0.25 | 0.25 | 4975 |
| **AE006468.2** | *Salmonella enterica* | Proteobacteria | 4,857,450 | 0.24 | 0.24 | 0.26 | 0.26 | 4453 |
| **AL590842.1** | *Yersinia pestis* | Proteobacteria | 4,653,728 | 0.26 | 0.26 | 0.24 | 0.24 | 4034 |

***Table1S.*** *Selection of five bacteria species and 8 genomic features up to 132 obtained using gbap.py. Size is the length of the entire genome, whereas fr_gen_A, fr_gen_T, fr_genC and fr_gen_G are the frequency of nucleotides divided by genome size. Fr_gen_A is the frequency of Adenine, fr_gen_T is the frequency of Thymine, fr_gen_C is the frequency of Cytosine, and fr_gen_G is the frequency of Guanine. N_cds is instead the number of protein coding genes (CDS) for each bacteria species.*

As an example of the potential of GBRAP we downloaded the GenBank files of some of the most studied bacterial species in the field of human health and food science: *Clostridium botulinum, Listeria monocytogenes* (Firmicutes class), *Escherichia coli, Salmonella enterica, and Yersinia pestis* (Proteobacteria class), in order to analyze the differences in their genomes. As shown in Figure1S, the percentage of A-T bases compared to C-G bases is strongly imbalanced in the Firmicutes group, whereas there is a certain equilibrium in the Proteobacteria class. These results are confirmed when graphing the boxplots reported in Figure2S, where we illustrated the existing differences in CDS Chargaff's second parity rule score (CT), together with Shannon's entropy. Our analysis demonstrated that, according to these two energetic constrains measures, the genomic structural stability is significantly higher in Proteobacteria compared to Firmicutes (Fariselli et al., 2021). Furthermore, following the same theory, we were able to state that the maximum value of both scores reached in the Proteobacteria class, confirms their higher content of genomic information, which is also underlined by their value of topological entropy and LZSS compression. Thanks to our tool, it is now possible to investigate the genome structural proprieties of different microorganisms using GBFF files. Looking exclusively to our results, the user can easily compare different level of genomic information, and eventually investigate the energetic proprieties of the DNA double helix of different microorganism species.

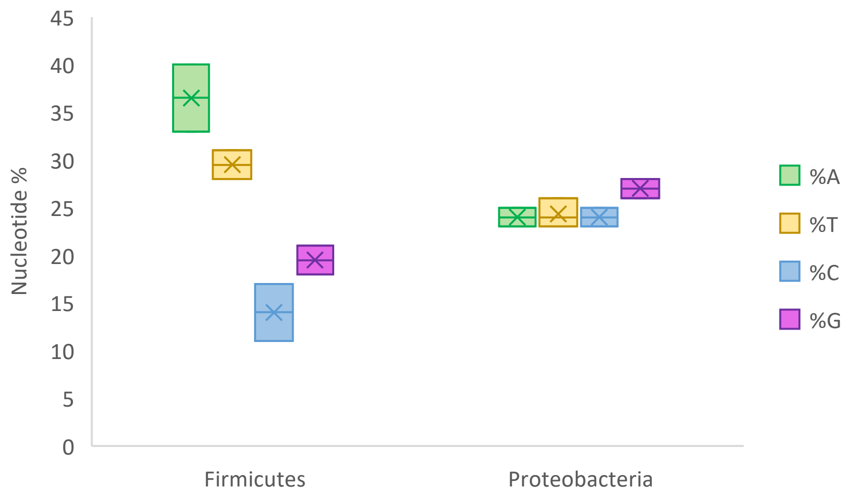


***Fig1S.*** *The percentage of nucleotides in the species of Firmicutes (C. Botulinum and L. monocytogenes) analyzed is more imbalanced compared to Proteobacteria organisms examined (E. Coli, S. enterica, and Y. pestis).*

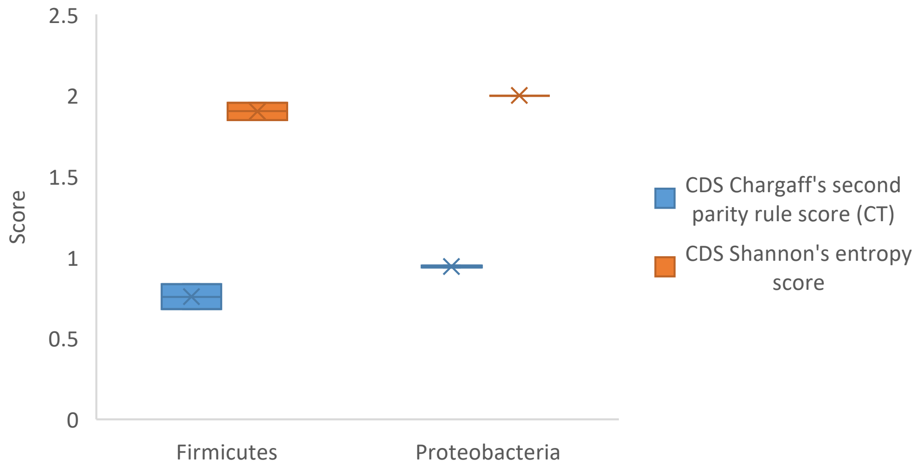


***Fig2S.*** *Shannon entropy and Chargaff's second parity rule (CT) score calculated for CDS (Coding Sequences) is lower in the species of Firmicutes (C. Botulinum and L. monocytogenes) analyzed compared to Proteobacteria organisms examined (E. Coli, S. enterica, and Y. pestis).*
